# Detecting selection with a genetic cross

**DOI:** 10.1101/2020.08.02.233106

**Authors:** Hunter B. Fraser

## Abstract

Distinguishing which traits have evolved under natural selection, as opposed to neutral evolution, is a major goal of evolutionary biology. Several tests have been proposed to accomplish this, but these either rely on false assumptions or suffer from low power. Here, I introduce a new approach to detecting lineage-specific selection that makes minimal assumptions and only requires phenotypic data from ∼10 individuals. The test compares the phenotypic difference between two populations to what would be expected by chance under neutral evolution, which can be estimated from the phenotypic distribution of an F_2_ cross between those populations. Simulations show that the test is robust to parameters such as the number of loci affecting the trait, the distribution of locus effect sizes, heritability, dominance, and epistasis. Comparing its performance to the QTL sign test—an existing test of selection that requires both genotype and phenotype data—the new test achieves comparable power with 50- to 100-fold fewer individuals (and no genotype data). Applying the test to empirical data spanning over a century shows strong directional selection in many crops, as well as on naturally selected traits such as head shape in Hawaiian *Drosophila* and skin color in humans. Applied to gene expression data, the test reveals that the strength of stabilizing selection acting on mRNA levels in a species is strongly associated with that species’ effective population size. In sum, this test is applicable to phenotypic data from almost any genetic cross, allowing selection to be detected more easily and powerfully than previously possible.

**Significance Statement:** Natural selection is the force that underlies the spectacular adaptations of all organisms to their environments. However, not all traits are under selection; a key question is which traits have been shaped by selection, as opposed to the random drift of neutral traits. Here, I develop a test of selection on quantitative traits that can be applied to almost any genetic cross between divergent populations or species. The test is robust to a wide range of potential confounders, and has greater power to detect selection than existing tests. Applied to empirical data, the test reveals widespread selection in both domesticated and wild species, allowing selection to be detected more easily and powerfully than previously possible.

## Introduction

Trait-based tests of selection aim to distinguish the effects of two major forces of evolution: natural selection and neutral drift. Because many factors affect trait divergence—e.g. population size, divergence time, and genetic architecture—distinguishing these two forces is seldom straightforward. Several types of trait-based selection tests have been proposed, all of which view neutrality as a null model, but which differ in how they assess this null and in the type of data they require (reviewed in Chapter 12 of Walsh and Lynch (1)).

For example, time series tests use phenotypic measurements of a single species over time, typically from the fossil record (a stratophenetic series). If the trait shows departure from the neutral expectation of a random walk—e.g. many more time steps with trait increases than decreases—then neutrality is rejected. The key assumption is that environmental changes do not affect these phenotypic trends, which is difficult to justify considering how much environments can change over the millions of years typically covered in a stratophenetic series.

A more widely used approach is known as Q_ST_, where the population structure of phenotypic variance is compared to the analogous genetic metric F_ST_. By utilizing genetic crosses in common garden experiments, the confounding effects of environment can be controlled, allowing selection to be assessed in a wide range of species (2). Limitations of this approach include low power (requiring data from >10 populations (3)) and several assumptions about epistasis and mutation rates (see Supplemental Note). However, an improved Q_ST_-based method has sufficient power to detect selection using only a few populations (4, 5).

Another widely used test is known as the quantitative trait locus (QTL) sign test (6, 7). In this test, QTL are first mapped using genotype and phenotype data from a genetic cross between two divergent parental lines. Under neutrality (and the absence of ascertainment bias), QTL directionality—i.e. which parent’s allele increases the trait at each QTL—is expected to be binomially distributed around 50%, much like a series of coin flips (Fig. 1a left). In contrast, under lineage-specific selection, QTL directions will be biased in one direction (Fig. 1a right). Although this test is quite robust due to its minimal assumptions, it also suffers from low power: a minimum of eight QTL (which is rarely reached in practice; see Supplemental Note) is required to achieve a nominal p < 0.01.

**Figure 1.**
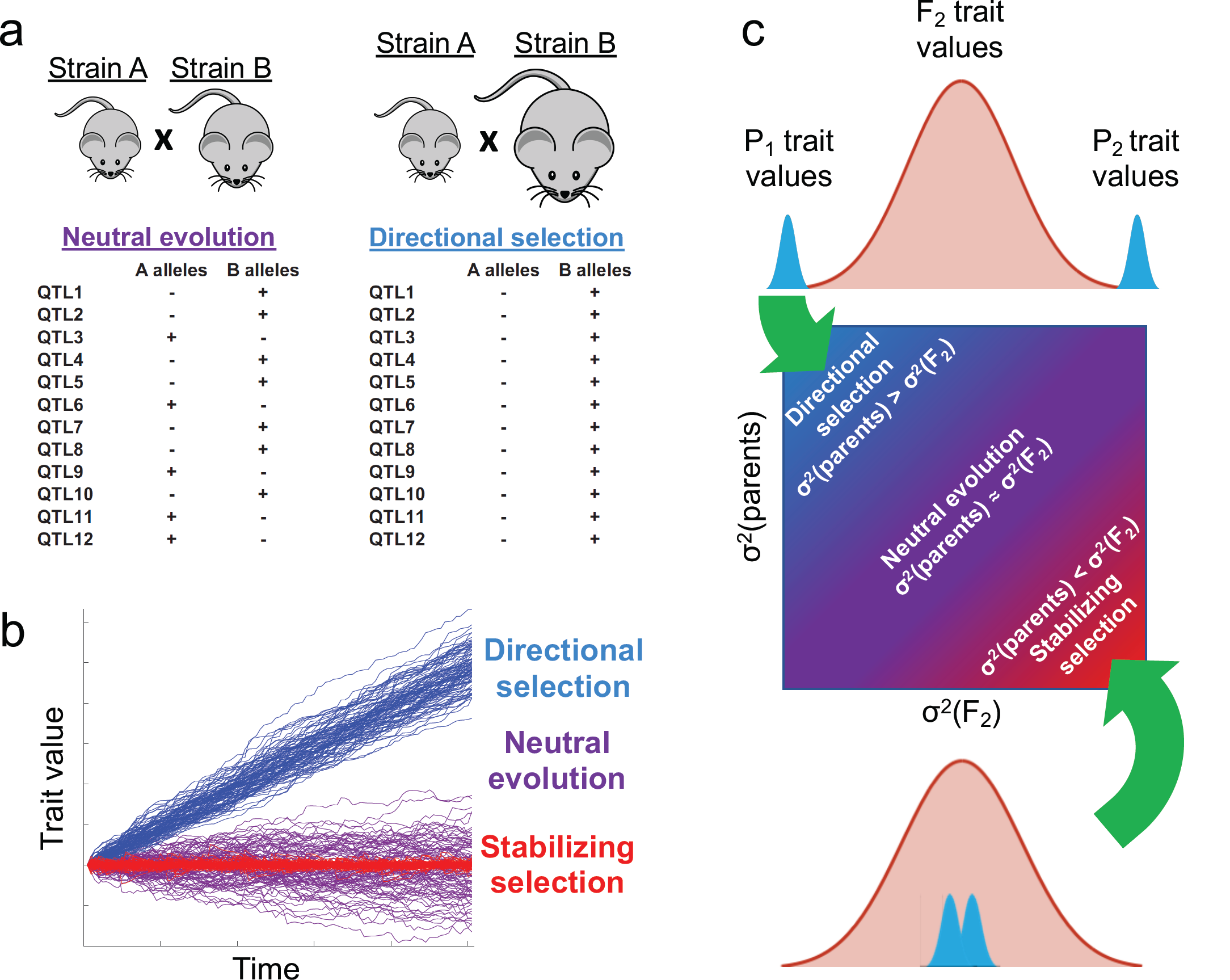
The sign test and the *v-*test. **a**. Illustration of the sign test applied to the trait of mouse size. Left panel: two mice from separate populations that have had no selection acting on size are expected to have approximately equal numbers of QTL (or QTN) alleles increasing size (binomially distributed with expected frequency = ½; stabilizing selection on size would result in a similar pattern, but with a smaller expected parental trait divergence). Right panel: In contrast, two populations that have experienced lineage-specific directional selection on size will show greater phenotypic divergence and a preponderance of QTL alleles increasing size in the larger strain. A significant deviation from the binomial expectation indicates rejection of the null hypothesis of neutral evolution. **b**. Simulation of trait divergence under a simple model of three selection regimes. One exponentially distributed QTL (or QTN) is added per time step and the number and effect sizes of QTL are identical in each selection regime; the only difference is their directionality. Under directional selection all QTL increase the trait value (as in Fig. 1a right panel); under neutral evolution their directionalities are random; and under stabilizing selection, their directionalities are whatever will bring the trait closer to the optimum (e.g. if the trait is above the optimum, the next QTL will be negative). Each selection regime has 100 lineages simulated for 100 time steps. **c**. Illustration of the *v*-test. Under a simple model, the variance of a neutral trait in two populations is expected to be approximately equal to that of their F_2_ progeny (Equation 1). Lineage-specific directional selection will result in higher parental variance, whereas stabilizing selection will lead to lower parental variance (transgressive segregation).

The sign test’s low power is largely due to the fact that it only uses QTL directionality information, while ignoring the phenotypic divergence between the two parental lines. However, the parental traits contain important information: if a trait evolves under directional selection, it will diverge much faster than under neutrality (Fig. 1b). If it were possible to estimate the divergence expected by chance under neutrality, then this could be used as a null hypothesis; parental trait divergence significantly greater than this expectation would suggest lineage-specific directional selection, whereas divergence less than this would suggest stabilizing selection.

Indeed, this intuitive logic underlies another class of trait-based methods, “rate tests,” that ask whether the phenotypic divergence of multiple populations is consistent with neutral drift (1, 8). The neutral expectation is estimated from population genetic theory, using parameters such as the effective population size, the mutational variance, and the number of generations since population divergence. Since these parameters and their sampling variances can typically only be roughly estimated (at best), and several strong assumptions must also be made, rate tests are viewed as qualitative guides rather than quantitative tests of neutrality (1, 8) (Supplemental Note).

In this work, I sought to develop a trait-based test of selection with the robustness of the sign test, while utilizing the framework of rate tests to increase the power to detect selection.

## Results

The logic underlying rate tests could lead to a more rigorous test of selection if the expected divergence under neutrality could be more accurately estimated. This can be achieved using the neutral model of the sign test, where the genetic variants underlying QTL (quantitative trait nucleotides, or QTN) have no directionality bias (Fig 1a left and Fig 1b purple). How could the distribution of phenotypes expected under this model of neutrality be estimated? One way would be to measure the effect size of every QTN and then predict the phenotypes resulting from random allelic combinations. However, there is a simpler solution: the F_2_ trait distribution represents exactly this null model. Regardless of the QTN directionalities in the parents, the F_2_ phenotypes result from random combinations of the segregating alleles; this randomness mimics the random directionality expected under neutral evolution (Fig 1a left, Fig 1b purple). Therefore every F_2_ individual can be thought of as a random draw from the distribution of potential parental phenotypes resulting from neutral evolution. If the parental divergence is significantly greater than expected based on the phenotypic variance observed in the F_2_ population, then neutrality can be rejected in favor of directional selection (Fig 1c top). If instead the parents are significantly less diverged than expected—known as transgressive segregation—then stabilizing selection is inferred (Fig 1c bottom).

I will begin with a simple model of a trait in a haploid species where all QTN are additive with equal effect sizes, and there is no environmental variation or trait measurement error (i.e. broad-sense heritability H^2^ = 1). Let *n*_*q*_ denote the number of QTN for this trait that differ between two strains/populations. Under neutrality, traits diverge from the ancestor like a random walk, proportional to 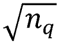 (Fig 1b purple). For two lineages evolving independently, the expected absolute difference will be 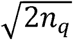. The difference between the two parental trait values represents a single draw from a binomial distribution with p = 0.5 and n = n_q_ (it is only one draw since once one parent’s allelic states are defined, the other’s must be the opposite for any QTN segregating in the cross); for n_q_ > ∼20 this approximates a normal distribution. The square of this difference is proportional to the parental trait variance; dividing this variance by the variance expected by chance under neutral evolution—i.e. the variance of the F_2_ trait distribution, as discussed above—results in the test statistic (denoted *v* and illustrated in Fig 1c):

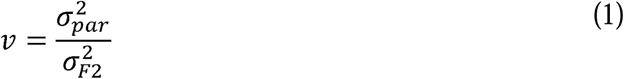

where 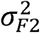 is the F_2_ phenotypic variance and 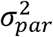 is the between-strain variance of the two parental strain (or population) means. This ratio is expected to be distributed as F(1, n_F2_-1) under neutrality, where n_F2_ is the number of F_2_ individuals. This approximates a χ^2^ distribution with 1 degree of freedom for n_F2_ > ∼20.

We can now relax the simplifying assumptions above. In diploids, the expected variance in the F_2_ is half that between the parents; this is accounted for by multiplying the denominator by a constant, denoted *c* (more generally, any factors that affect the phenotypic variance in the progeny—including dominance and other cross designs such as backcrosses or recombinant inbred lines [RILs]—can be accommodated by adjusting the value of this constant; see Supplemental Note). Allowing environmental variation and trait measurement error is equivalent to adding random noise to both the numerator and denominator. Correcting for this yields:

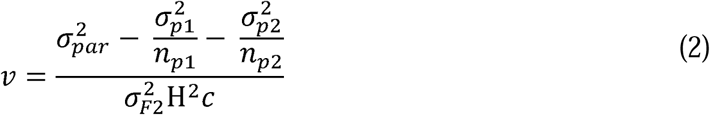

where *n*_*p*1_ and *n*_*p*2_ are the number of replicate individuals measured for each parental strain, and 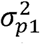 and 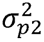 are the within-strain variances of each parental strain (see Supplemental Note). All of these terms can be estimated from phenotype data in a single genetic cross, provided that multiple individuals of each parental type are included. Note that Equation 1 is a special case of Equation 2, where c = 1 (for haploids) and H^2^ = 1 (hence 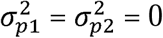).

To explore the behavior of *v* as a neutral null model, I conducted simulations of neutral traits in parental strains and their F_2_ progeny (see Methods). These simulations allowed the precise manipulation of individual parameters to assess their effects on *v*. These are presented as quantile-quantile (QQ) plots, where points following the Y=X line represent adherence to the expected F distribution. Points above and below the line represent values of *v* greater and less than expected under the null, respectively.

The test followed the expected distribution of *v* closely for a range of different QTL effect size distributions (Fig 2a). This includes the exponential distribution, which is thought to be a reasonable approximation for QTL (9, 10). However, extremely skewed distributions—e.g. a monogenic trait where one QTL explains all trait variance—will lead to half of all F_2_ individuals identical to one of the parents and *v* values tightly distributed around 1, resulting in little power to detect any deviation from the null. Therefore the test is only useful for polygenic traits.

**Figure 2.**
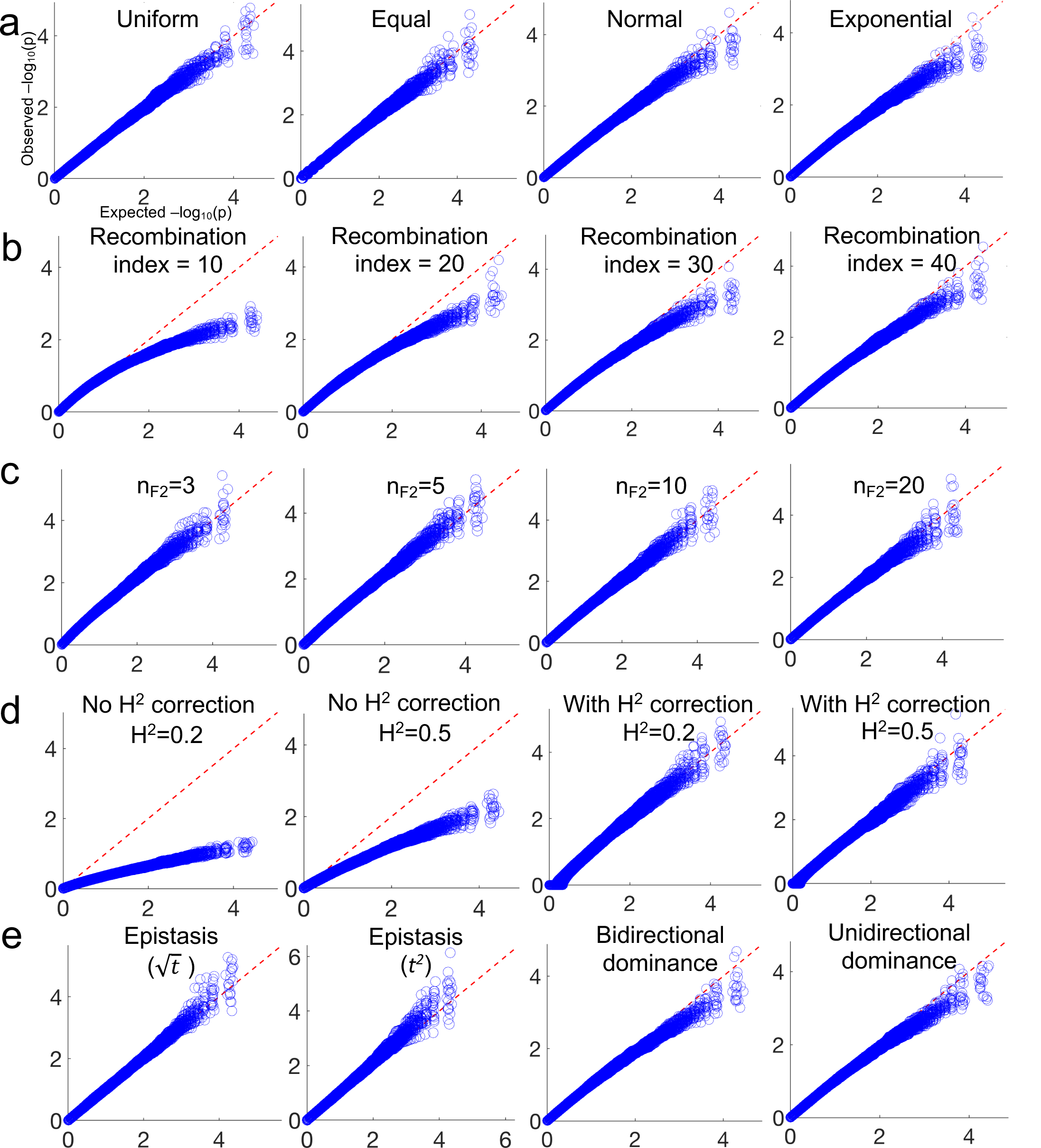
Neutral simulations. Each panel shows 20 quantile-quantile (QQ) plots where every point is an independent simulation of a genetic cross between two lineages where the trait in question has been evolving neutrally (i.e. QTL directions in each parent are random; Fig 1a-b). The X-axis shows expected p-value quantiles (uniform between zero and one), and the Y axis shows observed values of *v* (Equation 2) in 20 QQ plots each with 10^4^ simulations. For each panel, one parameter is varied from the baseline model (Exponential distribution of QTL effect sizes, recombination index = 50, number of parental replicates = 10, n_F2_= 100, H^2^=1, diploid, no epistasis or dominance), except for the upper right panel which is the baseline model. **a**. Effects of varying QTL effect sizes. **b**. Effects of varying recombination index. **c**. Effects of varying the number of F_2_ individuals. **d**. Effects of varying H^2^, with or without the correction in Equation 2. **e**. Effects of varying epistasis and dominance. Bidirectional dominance means all loci are fully dominant but with ∼50% of loci being dominant towards one parent, and ∼50% towards the other. Unidirectional means all loci are fully dominant in the same direction (i.e. the F_1_ phenotype is identical to one of the parents).

How polygenic must a trait be? More important than the number of genetic variants affecting the trait—which is likely to be quite large for complex traits (11)—is the number of independently segregating QTN, which is limited by the recombination index (12) (RI). The RI is defined as the haploid number of chromosomes plus the number of recombinations per meiosis; it represents the number of independent genomic regions segregating in a cross. Adherence of *v* to the null improves with greater RI (Fig 2b) since more shuffling of the two parental genomes leads to more normally distributed 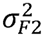 values. For example, an extreme case of a single non-recombining chromosome (RI = 1) would be equivalent to the monogenic example above where *v* cannot deviate far from 1. The *v*-test behaves conservatively in crosses with RI < ∼20 (Fig 2b). Fortunately, this is rarely an issue in practice since the mean RI for plants is ∼30 and for animals is ∼40 (13). (For RILs, there are more generations for recombination so the mean “effective RI” is ∼45 for plants and ∼58 for animals; see Methods.)

Sample size is another important consideration. Although more samples are always preferable, *v* follows the null distribution even with only three F_2_ individuals (Fig 2c).

Other potential sources of noise include environmental variability and measurement error. Although these affect both parental and F_2_ phenotypic variances, they have less effect on the parental estimates when parental replicates are included, because taking the mean of each parental strain reduces this noise. Without correcting for this effect (i.e. setting H^2^ = 1, and thus 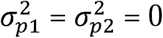, in Equation 2), low H^2^ leads to severe underestimates of *v* (Fig 2d left panels). However including a correction for this (Equation 2) precisely accounts for this effect (Fig 2d right panels).

The final effect explored via simulation is genetic interaction, which is the context-dependence of phenotypic effects either between loci (epistasis) or between alleles at the same locus (dominance). Epistasis can take many forms, but the large-scale pattern most often observed is known as diminishing returns epistasis (14–16), where strains with the greatest trait value have lower values than expected from additivity. To model this, I transformed the simulated trait values as 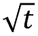, where *t* is each trait value. Since this affects both 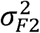 and 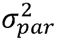 similarly, it has little impact on the distribution of *v* (Fig 2e). I also modeled synergistic (increasing returns) epistasis as *t*^2^, and again found no effect (Fig 2e). However, epistasis in more extreme forms could obscure any signal of selection (Supplemental Note). Dominance can be accounted for by adjusting the value of *c* to offset its effect on 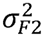 (Fig 2e, Supplemental Note).

In sum, simulations of neutral evolution show that *v* is robust with respect to the number of loci affecting the trait, the distribution of locus effect sizes, environmental variability, measurement error, dominance, and epistasis.

Having established the behavior of *v* under neutrality, I explored its power by simulating directional selection. These were identical to the neutral simulations except that all QTN had concordant parental directionality (Fig 1a right, Fig 1b blue). I compared the *v*-test to the sign test to assess their relative performance in a range of parameter settings (other selection tests use very different types of data, precluding a direct comparison). Both tests utilized the same simulated F_2_ phenotype data for each cross; the sign test was also provided with optimal genotype data, meaning that each QTN was genotyped without error, had no linkage with other QTN, and no other genetic markers were included. Both tests result in a probability of neutrality (p_nut_) value for each trait (see Methods). More extreme p_nut_ values represent greater power to detect selection, with points above the diagonal representing crosses with greater power for the *v*-test and those below the diagonal indicating greater power for the sign test.

Even with optimal genotype data, at small sample sizes (n_F2_ = 10 or 100) very few QTL are mapped, resulting in the sign test’s low power (Fig 3). In contrast, the *v*-test often rejects the null with p_nut_ < 10^−4^ even at n_F2_ = 10. The *v*-test is generally more powerful than the sign test at n_F2_ < 10^3^. However, the sign test generally outperforms the *v*-test at very large sample sizes (n_F2_ > 10^3^ when H^2^ > 0.8, and n_F2_ > 10^4^ for all H^2^).

**Figure 3.**
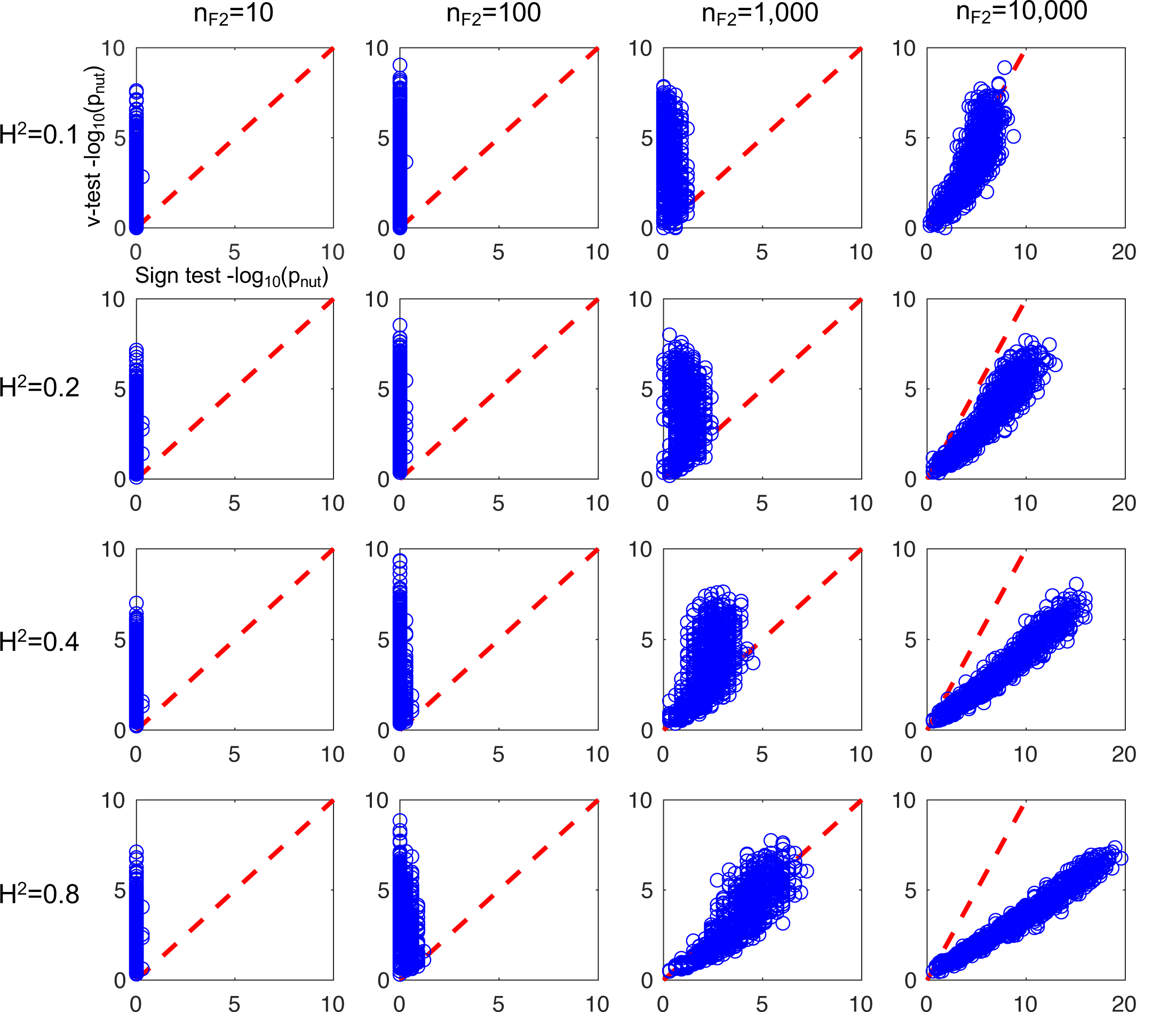
Directional selection simulations. All panels show scatter plots where every point is an independent simulation of a genetic cross between two lineages where the trait in question has been evolving under directional selection (i.e. all QTL are in the same direction; Fig 1a-b). The X-axis shows sign test log p-values, and the Y axis shows *v* test log p-values. For each panel, two key parameters (H^2^ and n_F2_) are set to the values shown and a third is varied within the panel (RI, which takes on all integer values from 5 to 100). For each value of RI, 10 simulations are shown, each with an independent set of QTL effect sizes; this results in 960 simulations (data points) per panel. All other parameters are kept constant throughout the figure (Exponential distribution of QTL effect sizes, n_par_ = 10, diploid, no epistasis or dominance).

To further explore the *v*-test’s power at small sample sizes, I simulated crosses with a total of 10, 20, or 30 phenotyped individuals (including the parents). For example at n = 10, 93% of traits rejected the null at p_nut_ < 0.05 when H^2^ = 0.1, and 99% when H^2^ = 0.8 (Supp Fig 1a). The *v*-test performed well with 10 individuals even when selection was weak (with up to 20% of QTN acting in opposition to the majority) and heritability was low (Supp Fig 2). In contrast, the sign test required 500-1000 phenotyped and genotyped individuals to reach the same power (Supp Fig 1b). This difference in power of the two tests makes sense considering that although they are both evaluating the same neutral null model, the sign test is doing so directly (by comparing QTL directionalities to the neutral expectation; Fig 1a), whereas the *v*-test is doing so indirectly. When enough QTL are mapped, the direct approach of the sign test is superior, but by not needing to map QTL, the *v*-test requires fewer individuals, as well as no genotype data.

Due to its generality, the *v*-test can be applied to data from almost any genetic cross where both parental strains and their F_2_ (or other cross) progeny were phenotyped. This includes most QTL studies, as well as genetic studies from before genotyping was possible. To explore the test empirically, I collected published data for 126 traits from 21 species (Supp Table 1). The *v*-test p_nut_-values for artificially selected traits (in crops, livestock, and laboratory experiments) revealed a strong skew towards low p_nut_-values indicating directional selection, as expected(17) (Fig 4a; for a discussion of trait ascertainment bias—a major caveat for all trait-based selection tests—see Discussion and Supplemental Note). Three of the most significant traits were from maize, including data for ear length and seed weight published in 1913, suggesting intense artificial selection on these traits prior to this date (18).

**Figure 4.**
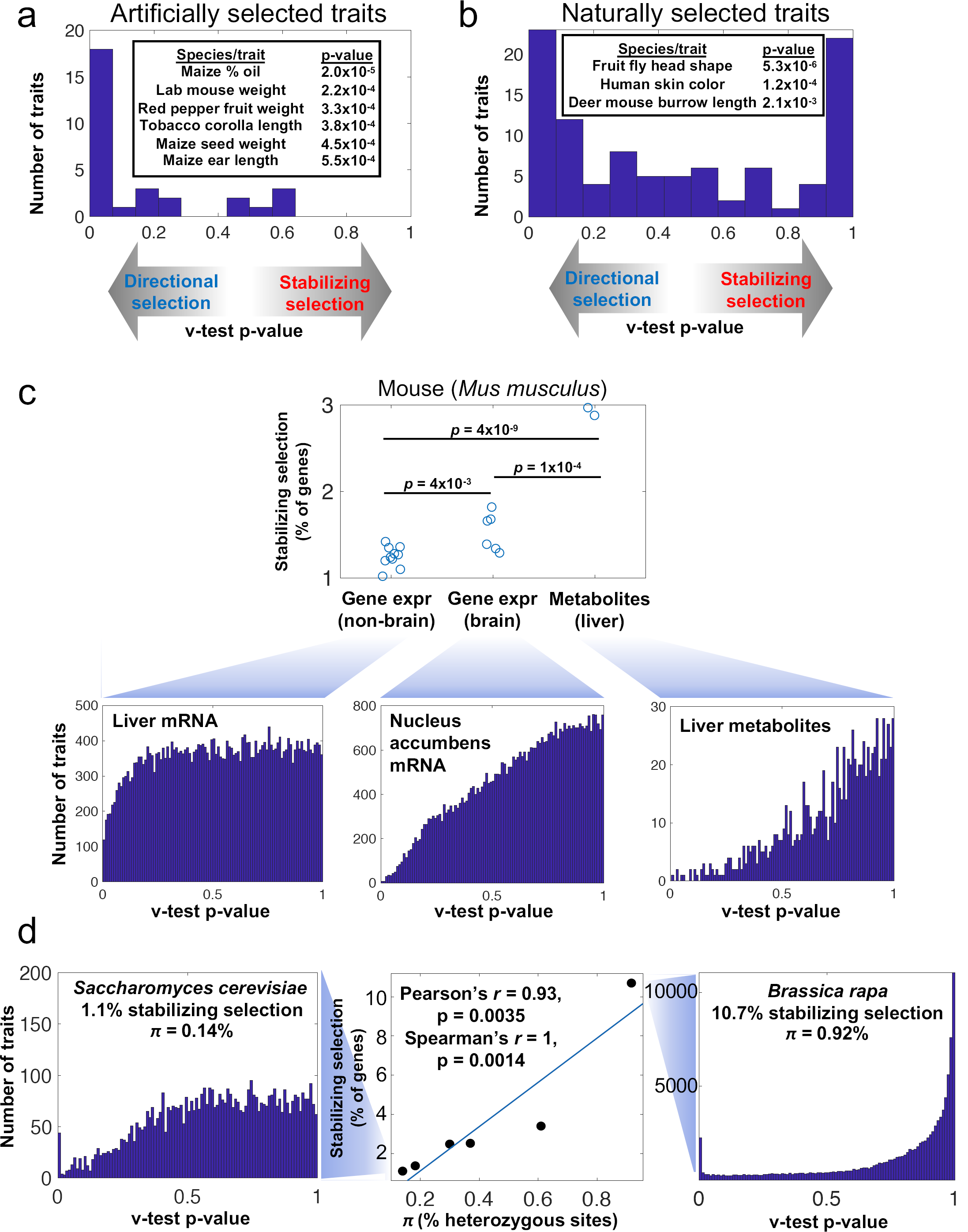
Empirical analysis. **a**. Results for artificially selected traits in crops, livestock, and laboratory selection experiments. Inset shows the six most significant traits. **b**. Results for naturally selected traits in plants and animals. Inset shows the three most significant traits. **c**. Results for gene expression (mRNA levels) and metabolite levels measured in the same mouse RIL panel (BXD). Note that any selection detected between the two parental lineages could involve divergence of their wild ancestors (mostly *M. musculus domesticus*) and/or artificial selection during their inbreeding in the lab. T-test p-values shown for each comparison. **d**. Center panel: The strength of stabilizing selection vs. heterozygosity (π) in six species (in order of decreasing π: *Brassica rapa, Arabidopsis thaliana, Caenorhabditis elegans, Oryza sativa, Mus musculus, Saccharomyces cerevisiae*). Side panels: the full distribution of p_nut_ values for the species with the highest (right) and lowest (left) π.

In contrast, the p_nut_-value distribution for traits naturally selected in the wild showed a less extreme skew (Fig 4b; Supp Table 1; comparing distributions, Wilcoxon p = 2×10^−5^), with peaks at both low and high p-values suggesting a wide range of selection pressures. The most significant trait for directional selection was male head shape in a cross between two species of Hawaiian *Drosophila* (19) (data from reciprocal F_2_ crosses and an F_6_ cross are all significant; Supplemental Note). This is a well-known example of rapid morphological evolution, potentially due to sexual selection (20), but whether the divergence could instead be explained by genetic drift was not previously testable. The next most significant trait was human skin color, measured in the “F_2_” grandchildren of West Africans and Europeans (21). Despite the small sample size (n_F2_ = 14), the *v*-test is significant at all three reflectance wavelengths tested (Supp Table 1). Human skin color has long been thought to be adaptive based on its correlation with local UV radiation (22); these results provide independent confirmation for the role of selection. The third most significant wild trait was burrowing behavior in *Peromyscus* mice (23), as measured by burrow length in an interspecies backcross.

The *v*-test can also be applied to molecular-level traits such as gene expression levels, which avoids the effects of trait ascertainment bias (Supp Note). The BXD collection of mouse RILs is an excellent test case, with gene expression data available for 16 tissues. Performing the *v*-test revealed that the p_nut_-value distributions of all tissues were shifted towards stabilizing selection to varying degrees (Fig 4c). To estimate the strength of stabilizing selection in each tissue, I calculated the fraction of genes with *v*-test p_nut_ > 0.99. All 16 tissues had values between 1.0%-1.8% (note this should not be interpreted as the fraction of genes under stabilizing selection, which is likely to be far higher). Interestingly, gene expression from six different regions of the brain had significantly stronger stabilizing selection than in the ten non-brain tissues (Fig 4c). This is consistent with previous reports of slower evolution of gene expression in the mammalian brain compared to other tissues(24, 25), and suggests that this slower evolution is at least partially due to greater selective constraint (as opposed to a lower mutational variance, which can also lead to a slower evolutionary rate (1)).

Another type of molecular trait measured in the BXD cross is metabolite levels in liver (26). Applying the *v*-test to these metabolomic data, cohorts fed two diets (high-fat and normal chow) showed p_nut_-values strongly skewed towards one (Fig 4c). Therefore the measured metabolites appear to be under stronger stabilizing selection than mRNA levels.

Despite the variation in stabilizing selection on gene expression across the 16 tissues, all tissues were relatively close to the neutral expectation (1% of genes at p_nut_ > 0.99). To compare this to other species, I collected gene expression data from genetic crosses of five additional species (*Saccharomyces cerevisiae, Oryza sativa, Arabidopsis thaliana, Brassica rapa*, and *Caenorhabditis elegans*; Supp File 1). In contrast to mouse, some species had much stronger stabilizing selection (e.g. *B. rapa* with 10.7% of genes under stabilizing selection). One hypothesis to explain this wide range of values is that natural selection is expected to be stronger in species with larger effective population sizes (N_e_). Direct measurements of N_e_ for these species are not possible, but an indirect indicator is the fraction of neutral genomic positions that are heterozygous (known as π), which is expected to increase with N_e_ (27). Plotting the strength of stabilizing selection against published values of π, I observed a strong correlation (Fig 4d). This suggests that N_e_ (or another factor correlated with N_e_; see Supp Note) may be a major determinant of stabilizing selection on gene expression levels, as has been previously proposed (24, 25).

The peaks of gene expression p_nut_-values near zero (Fig 4d) suggest that directional selection may also be detectable from these data. For example, in *S. cerevisiae* genes with low p_nut_ are highly enriched for roles in mitochondrial translation (FDR = 6×10^−23^ among genes down-regulated in the lab strain BY; no enrichment among genes up-regulated in BY). In *B. rapa*, the defense response to other organisms is the most enriched function among genes with low p_nut_ (FDR = 0.02). Similarly in *C. elegans*, immune response was the most enriched (FDR = 0.04).

## Discussion

In this work, I introduced a new test of selection that combines the logic of rate tests with the neutral null model of the sign test. The result is a test that is simple, robust, and more powerful than existing tests. This will allow researchers to assess selection on traits of interest any time they perform a genetic cross. Moreover, if multiple traits are measured in the same cross, results from the test can be directly compared to assess the strength of selection on diverse phenotypes (as in Fig 4c).

There are many potential extensions to this test. For example, the *v*-test framework could be applied any time the genomes of two divergent populations are mixed, including naturally admixed populations. In this case, the only modification needed would be in the calculation of c, representing the expected ratio of parental variance to admixed progeny variance (Supplemental Note). Other extensions could include testing multiple correlated traits simultaneously, focusing only on additive effects, or estimating confidence intervals (Supplemental Note).

It is important to note that results from all trait-based tests of selection must be treated with caution when trait ascertainment bias is present. If traits are chosen for study based on the extent of their divergence between populations, then the neutral model no longer holds. For example, imagine 100 neutral traits; in any properly calibrated selection test, we would expect ∼5 of these to reach a nominal p < 0.05. If these same five are the only traits included in a study (e.g. because they have the strongest phenotypic divergence), then they will appear to be inconsistent with whichever null model they are tested against. In some cases it is possible to correct for ascertainment bias, either by modifying the test itself (6) or by using a more conservative p-value threshold (see Supplemental Note). However, the ideal solution is to analyze traits that were selected for study independently of the parental trait values, which by definition lack any ascertainment bias. The most widespread examples of this are molecular-level traits such as the levels of mRNAs, proteins, metabolites, etc. Similarly, standardized phenotyping (28) can be free of bias.

Notably, Equation 2 is identical to the widely used Castle-Wright (CW) estimator for the number of loci underlying divergence in a quantitative trait (29–31) (though it was derived independently). The maximum possible value of this estimator is the RI of the species being crossed, resulting in a strong downward bias for most complex traits, which can have thousands of variants contributing (11). It is therefore rather fortuitous that this severely biased estimator is also precisely F-distributed under the null hypothesis of neutrality, even though neutrality had no role in its original derivation and the F distribution has no role in its traditional interpretation (29, 30). Furthermore, the true number of loci underlying a trait (what the CW estimator aims to estimate) is not indicative of selection; a neutral trait could have any number of underlying loci, so this cannot be used to assess a trait’s neutrality.

How can this one equation have two seemingly unrelated interpretations? The CW estimator requires a number of restrictive assumptions, including that all QTL must act in the same direction with respect to their parent of origin (as in Fig 1a right panel). Rather than being an assumption of the *v*-test, this concordant QTL directionality is exactly what the *v*-test was designed to detect. Therefore the connection between the two interpretations rests on the fact that the strength of the signal detected by the CW estimator—the number of reinforcing QTL acting in the same parental direction—is indicative not only of the number of loci, but also of any selection that has acted on those loci since the divergence of the two parental strains.

One consequence of this mathematical homology is that the hundreds of published CW estimator values dating back to 1921 (29) can now be immediately reinterpreted as tests of neutral evolution (Supplemental Note), even when the phenotype data from these studies are not available.

## Methods

### Neutral simulations

Parameters in neutral simulations are shown in Fig 2: QTN effect size distribution, RI (the number of independently segregating QTN), n_F2_, H^2^, epistasis, and dominance (c = 2 for additive, c = 1 for bidirectional dominant, and c = 4/3 for unidirectional dominant; see Supplemental Note). Parameter values were chosen to reflect typical published data sets rather than optimal parameters for test performance. First I generated effect sizes for the specified number of QTN by sampling from the specified distribution. In neutral evolution, QTN directions in each parent are random (Fig 1a), so the first parent’s traits were determined by flipping the sign of each effect size with 50% chance and then summing the values. The second parent’s traits were calculated the same way but with all signs flipped from the first parent (each QTN increases the parental difference by twice its effect size, assuming parents are homozygous). F_2_ traits were determined by multiplying each effect size by one number chosen randomly from a set of four numbers that represent the four possible F_2_ genotypes at each locus and their resulting phenotypic effects ([0 1 1 2] for additive, [0 2 2 2] for unidirectional dominant, or [0 0 2 2] for bidirectional dominant; see Supplemental Note), separately for every individual, and summing across all QTN. When H^2^ < 1, random noise was also added to each parental and F_2_ individual as a normally distributed variable with zero mean and standard deviation 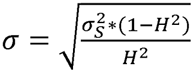, where 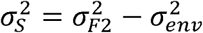 (which in practice is calculated as the variance of the F_2_ trait values before noise is added, since 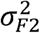 and 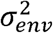 are not known until the noise is calculated). In epistasis simulations, traits were additionally transformed either by 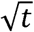 or *t*^2^, where *t* is the trait value. Parental within-strain variances (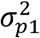 and 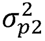) were then estimated from 10 replicates per parent, and 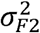 was estimated from the F_2_ population. From these variables, *v* was calculated using Equation 2 and converted into a *p*-value based on the F(1, n_F2_-1) cumulative distribution.

### Selection simulations

Selection was simulated using the framework described above, with one difference: omitting the step of flipping the sign of each effect size with 50% chance. This meant that all QTN were reinforcing in their directionalities. *v* was then calculated as described above and converted to a p-value based on the F(1, n_F2_-1) cumulative distribution. Parameter values are listed in Fig 3 legend.

To calculate the sign test p-value, QTL must first be mapped. The genotype of each QTN variant (randomly generated as described above) was provided in a genotype matrix, with no genotyping error and no additional genetic markers (this represents an unrealistic best-case scenario for QTL mapping). Pearson’s correlation coefficient between each marker and the F_2_ phenotypes were then converted into LOD scores (32) as 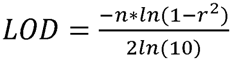. LOD > 3 was required to call a QTL. The directionalities for the full set of QTL for each cross were then assessed for their fit to the binomial distribution cumulative distribution with expected frequency = ½. The resulting two-sided p-value was the sign test p-value.

The simulations in Supp Figs 1-2 were identical to those in Fig 3, but with different parameter values and different visualizations. 1000 simulations were performed for each combination of parameter values shown. In Supp Fig 2, true positives were defined as crosses simulating selection where the *v*-test p-value was below a given cutoff; false positives were crosses simulating neutral evolution where the p-value was below the same cutoff (the cutoff varied from 0 to 1 to generate each ROC curve). Selection strength was represented by the fraction of QTN with reinforcing directionality. This cannot be translated into a selection coefficient since it would depend on how the trait relates to fitness, the population size, time since population divergence, and many other parameters.

### Empirical analysis

Traits were collected for Fig 4a from two sources: Wright (30) Tables 15.1-15.8, and Lynch and Walsh (32) Table 9.6. Traits were collected for Fig 4b from Lynch and Walsh (32) Table 9.6, Rieseberg (17) Table 5, and literature searches for cichlid and *Peromyscus* data.

For traits in Fig 4a-b and Supp Table 1, H^2^ values were estimated as follows. If values were provided by the authors of the original study, these were used. If not, the environmental variance was estimated using Wright’s preferred method (30), a weighted average of within-strain variances: 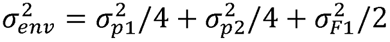. If data from the F_1_ were not available, then I used the sample size of each parental type as weights: 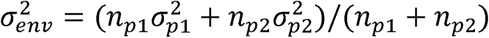. This variance was then used to calculate 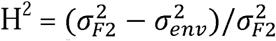. In some cases this can lead to a negative H^2^ (likely due to overestimation of 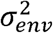 since this was always based on fewer replicates than 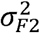); therefore values of H^2^ < 0.1 were set to 0.1. For the two cases of traits with outbred parents (burrowing and parenting behavior in *Peromyscus*), H may be underestimated due to within-strain genetic variation contributing to 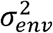; therefore I conservatively set H^2^ = 0.4 for these traits (which is higher than the heritability of most behavioral traits (33)), resulting in less significant values of p_nut_ (see Supplemental Note).

Data in Fig 4c were collected from http://www.genenetwork.org/ (selecting species: mouse; group: BXD family; type: any tissue with parental data). Since most mouse gene expression data sets in Fig 4c had only one sample per parental strain, H^2^, 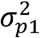, and 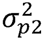 could not be accurately estimated. To allow a direct comparison between data sets with parental replicates vs. those without, I made two modifications: 1) for all tissues, I assumed half of the parental variance was genetic, and half environmental (i.e. the numerator of Equation 2 was set to 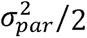). 2) I set H^2^=0.31 for all genes, this being the median value estimated for yeast (34) (specifically the median of 1-*e*, where *e* is the residual gene expression variance not explained by either additive or pairwise epistatic effects). I selected yeast because it had the largest number of gene expression profiles from a genetic cross of any species, a comprehensive heritability analysis performed by the original authors, and the closest π to mouse. Note that these modifications affect the Y-axis values in Fig 4c, but not the relative relationship between points; any values could have been used without affecting the trend shown. The complete list of tissues and stabilizing selection scores are in Supp File 1.

Gene expression data for the six species in Fig 4d were from the following sources: *S. cerevisiae* (34), *A. thaliana* (35), *B. rapa* (36) (normal phosphorus condition), *C. elegans* (37, 38) (control condition), *O. sativa* (39), and *M. musculus* (see above). Published expression data from other species’ crosses were not usable (e.g. no parental data). For mouse, the median stabilizing selection level across all 16 tissues was used. To avoid spuriously low or negative estimates of H^2^, for all six species any genes with H^2^ < 0.1 were set to 0.1 (as described above). As above, to facilitate comparison across data sets I assumed half of the parental variance was genetic, and half environmental. All p_nut_ values are listed in Supp File 1. π values were taken from Leffler et al. (40) as the median of autosomal π estimates for each species. No values were listed for *O. sativa* or *B. rapa*, so other published estimates were used (41, 42). For *O. sativa*, both parents of the genetic cross (Zhengshan 97 and Minghui 63) were in population group XI, so the π for this group was used. Using a more recently published π estimate for *S. cerevisiae* (0.18%, which is the median π across all 14 non-mosaic populations (43)) yielded a slightly stronger Pearson correlation (*r* = 0.935). Omitting *B. rapa* as an outlier also strengthened the correlation (*r* = 0.960). Gene Ontology process enrichments were calculated with GOrilla (44).

### Estimating recombination index

RI was estimated as [mean cM/100] + [mean haploid chromosome number] for 189 plants and 140 animals (13). RILs experience around twice as many detectable recombinations (defined as those occurring in a heterozygous genomic region) as an F_2_ cross, regardless of the exact number of generations of inbreeding, so for RILs the recombination values were doubled. Backcrosses experience about half as many detectable recombinations as an F_2_ cross.

## Acknowledgements

I would like to thank M. Feldman, A. Kern, D. Petrov, M. Schumer, and the Fraser lab for helpful discussions and feedback, and C. Hu and J. Bloom for sharing data. HBF is supported by NIH grant 1R01GM13422801.

---

**Supplemental Figure 1.**
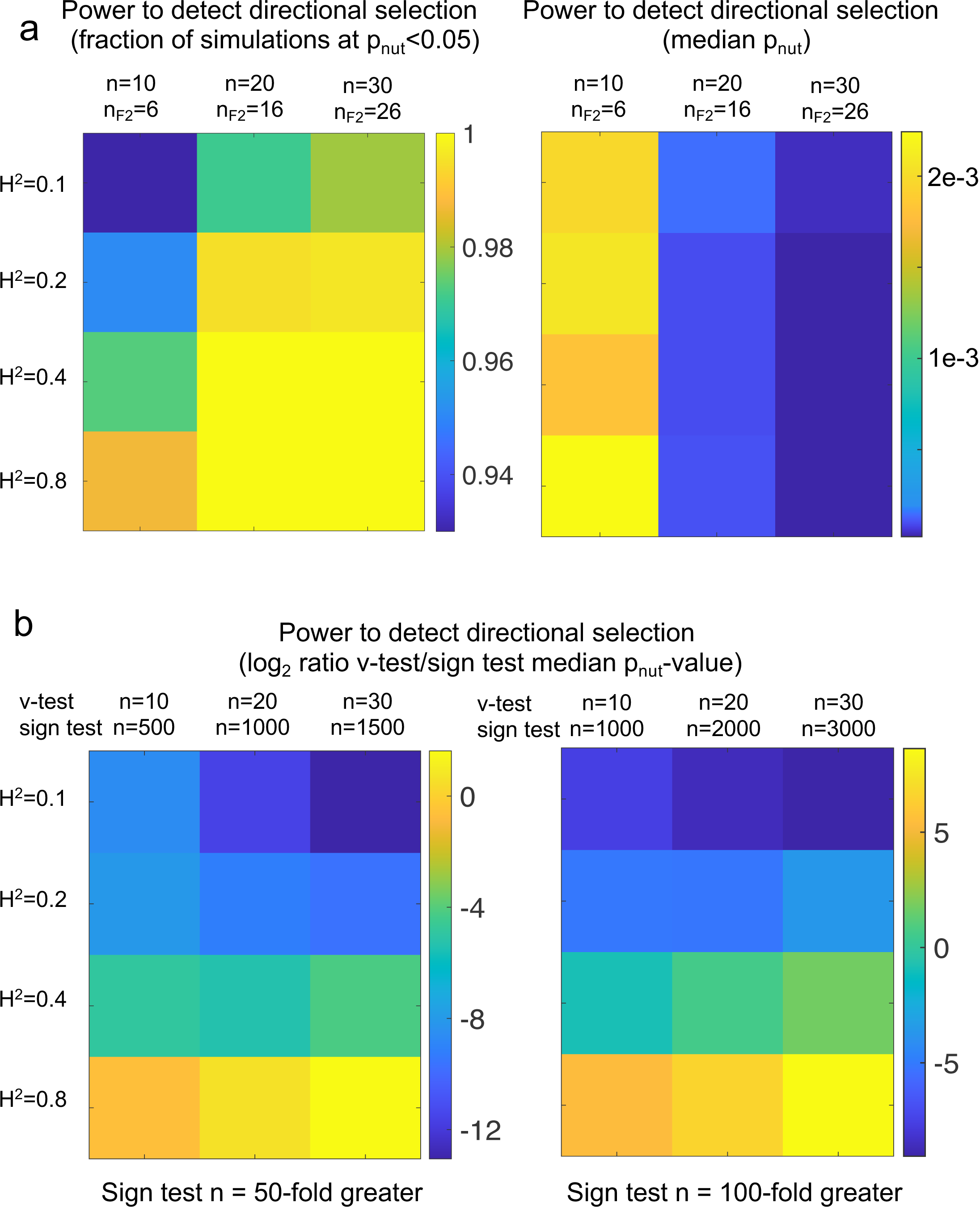
**a**. Directional selection was simulated as in Figure 3. Two replicates of each parental line were used together with the indicated number of F_2_ individuals. Therefore the total number of individuals n = n_F2_ + 4. **b**. The *v*-test simulations from panel (a) were compared to sign test results using 50-fold (left) or 100-fold (right) more individuals (all F_2_ since the sign test does not require parental data). Negative values indicate the *v*-test had lower median p-value than the sign test in that comparison. The sign test generally requires >50-fold more phenotyped individuals (as well as genotypes) to reach the same median p-value as the *v*-test; for traits with low H^2^ it requires >100-fold more individuals.

**Supplemental Figure 2.**
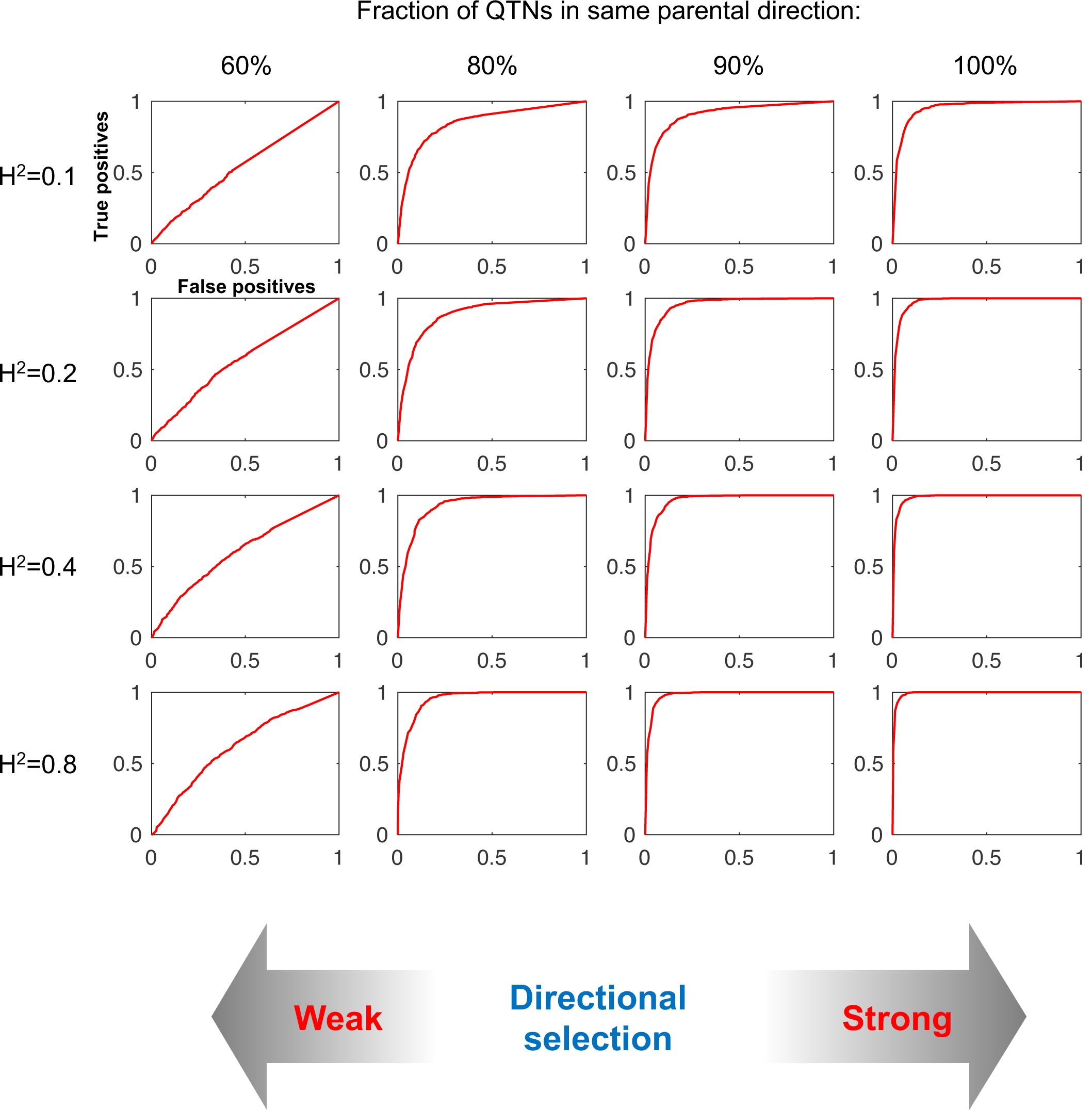
Receiver-operator characteristic curves are shown for the *v*-test with a range of H^2^ and selection strengths. In these plots, a perfect classifier would have 100% true positives and 0% false positives; a random classifier would be on the diagonal X=Y line. All simulations used 10 individuals (two of each parental strain and six F_2_), exponential distribution of QTN effect sizes, RI = 50, diploid, and no epistasis or dominance. Neutral evolution was simulated as in Figure 2 (defined as having QTN directionality binomially distributed around 50%). Directional selection was simulated as in Figure 3, except that the fraction of reinforcing QTN (reflecting the strength of selection) was allowed to vary. Note that in these simulations QTN directionality was independent of effect size; a more realistic case is that QTN of larger effect would be less likely to oppose the direction of selection, in which case the test performance would increase for any given % of reinforcing QTN.

## References

1. B. Walsh, M. Lynch, Evolution and Selection of Quantitative Traits (2018).

2. T. Leinonen, R. B. O’Hara, J. M. Cano, J. Merilä, Comparative studies of quantitative trait and neutral marker divergence: A meta-analysis. J. Evol. Biol. 21, 1–17 (2008).

3. R. B. O’Hara, J. Merilä, Bias and precision in QST estimates: Problems and some solutions. Genetics 171, 1331–1339 (2005).

4. R. B. O’Hara, et al., driftsel: An R package for detecting signals of natural selection in quantitative traits. Mol. Ecol. Resour. 13, 746–754 (2013).

5. O. Ovaskainen, M. Karhunen, C. Zheng, J. M. C. Arias, J. Merilä, A new method to uncover signatures of divergent and stabilizing selection in quantitative traits. Genetics 189, 621–632 (2011).

6. H. A. Orr, Testing natural selection vs. genetic drift in phenotypic evolution using quantitative trait locus data. Genetics 149, 2099–104 (1998).

7. H. B. Fraser, Genome-wide approaches to the study of adaptive gene expression evolution. BioEssays 33, 469–477 (2011).

8. M. Turelli, J. H. Gillespie, R. Lande, RATE TESTS FOR SELECTION ON QUANTITATIVE CHARACTERS DURING MACROEVOLUTION AND MICROEVOLUTION. Evolution (N. Y). 42, 1085–1089 (1988).

9. S. P. Otto, C. D. Jones, Detecting the undetected: estimating the total number of loci underlying a quantitative trait. Genetics 156, 2093–107 (2000).

10. H. A. Orr, THE POPULATION GENETICS OF ADAPTATION: THE DISTRIBUTION OF FACTORS FIXED DURING ADAPTIVE EVOLUTION. Evolution (N. Y). 52, 935–949 (1998).

11. E. A. Boyle, Y. I. Li, J. K. Pritchard, An Expanded View of Complex Traits: From Polygenic to Omnigenic. Cell 169, 1177–1186 (2017).

12. C. D. Darlington, The Biology of Crossing-over. Nature 140, 759–761 (1937).

13. J. Stapley, P. G. D. Feulner, S. E. Johnston, A. W. Santure, C. M. Smadja, Variation in recombination frequency and distribution across eukaryotes: patterns and processes. Philos. Trans. R. Soc. B Biol. Sci. 372, 20160455 (2017).

14. S. Kryazhimskiy, D. P. Rice, E. R. Jerison, M. M. Desai, Microbial evolution. Global epistasis makes adaptation predictable despite sequence-level stochasticity. Science 344, 1519–1522 (2014).

15. H.-H. Chou, H.-C. Chiu, N. F. Delaney, D. Segrè, C. J. Marx, Diminishing returns epistasis among beneficial mutations decelerates adaptation. Science 332, 1190–2 (2011).

16. A. I. Khan, D. M. Dinh, D. Schneider, R. E. Lenski, T. F. Cooper, Negative epistasis between beneficial mutations in an evolving bacterial population. Science 332, 1193–6 (2011).

17. L. H. Rieseberg, A. Widmer, A. M. Arntz, J. M. Burke, Directional selection is the primary cause of phenotypic diversification. Proc. Natl. Acad. Sci. 99, 12242–12245 (2002).

18. R. A. Emerson, E. M. East, The inheritance of quantitative characters in maize. Res. Bull. Bull. Agric. Exp. Stn. Nebraska No.2 (1913).

19. A. R. Templeton, Analysis of Head Shape Differences Between Two Interfertile Species of Hawaiian Drosophila. Evolution (N. Y). 31, 630 (1977).

20. C. R. B. Boake, Sexual selection and speciation in Hawaiian Drosophila. Behav. Genet. 35, 297–303 (2005).

21. G. A. Harrison, J. J. Owen, STUDIES ON THE INHERITANCE OF HUMAN SKIN COLOUR. Ann. Hum. Genet. 28, 27–37 (1964).

22. N. G. Jablonski, G. Chaplin, The colours of humanity: the evolution of pigmentation in the human lineage. Philos. Trans. R. Soc. B Biol. Sci. 372, 20160349 (2017).

23. J. N. Weber, B. K. Peterson, H. E. Hoekstra, Discrete genetic modules are responsible for complex burrow evolution in Peromyscus mice. Nature 493, 402–405 (2013).

24. D. Brawand, et al., The evolution of gene expression levels in mammalian organs. Nature 478, 343–348 (2011).

25. P. Khaitovich, W. Enard, M. Lachmann, S. Pääbo, Evolution of primate gene expression. Nat. Rev. Genet. 7, 693–702 (2006).

26. E. G. Williams, et al., Systems proteomics of liver mitochondria function. Science (80-.). 352 (2016).

27. M. Kimura, The neutral theory of molecular evolution (1983).

28. T. F. Meehan, et al., Disease model discovery from 3,328 gene knockouts by The International Mouse Phenotyping Consortium. Nat. Genet. 49, 1231–1238 (2017).

29. W. E. Castle, AN IMPROVED METHOD OF ESTIMATING THE NUMBER OF GENETIC FACTORS CONCERNED IN CASES OF BLENDING INHERITANCE. Science (80-.). 54, 223–223 (1921).

30. S. Wright, Evolution and the Genetics of Populations, Volume 1 (1968).

31. C. C. Cockerham, Modifications in estimating the number of genes for a quantitative character. Genetics 114, 659–664 (1986).

32. M. Lynch, B. Walsh, Genetics and analysis of quantitative traits (1998).

33. D. G. Stirling, D. Réale, D. A. Roff, Selection, structure and the heritability of behaviour. J. Evol. Biol. 15, 277–289 (2002).

34. F. W. Albert, J. S. Bloom, J. Siegel, L. Day, L. Kruglyak, Genetics of trans-regulatory variation in gene expression. Elife 7, 1–39 (2018).

35. M. A. L. West, et al., Global eQTL Mapping Reveals the Complex Genetic Architecture of Transcript-Level Variation in Arabidopsis. Genetics 175, 1441–1450 (2007).

36. J. P. Hammond, et al., Regulatory Hotspots Are Associated with Plant Gene Expression under Varying Soil Phosphorus Supply in Brassica rapa. Plant Physiol. 156, 1230–1241 (2011).

37. M. G. Sterken, et al., Dissecting the eQTL micro-architecture in *Caenorhabditis elegans*. bioRxiv, 651885 (2019).

38. B. L. Snoek, et al., Contribution of trans regulatory eQTL to cryptic genetic variation in C. elegans. BMC Genomics 18, 500 (2017).

39. J. Wang, et al., An expression quantitative trait loci-guided co-expression analysis for constructing regulatory network using a rice recombinant inbred line population. J. Exp. Bot. 65, 1069–1079 (2014).

40. E. M. Leffler, et al., Revisiting an Old Riddle: What Determines Genetic Diversity Levels within Species? PLoS Biol. 10, e1001388 (2012).

41. S. Park, H.-J. Yu, J.-H. Mun, S.-C. Lee, Genome-wide discovery of DNA polymorphism in Brassica rapa. Mol. Genet. Genomics 283, 135–145 (2010).

42. W. Wang, et al., Genomic variation in 3,010 diverse accessions of Asian cultivated rice. Nature 557, 43–49 (2018).

43. J. Peter, et al., Genome evolution across 1,011 Saccharomyces cerevisiae isolates. Nature 556, 339–344 (2018).

44. E. Eden, R. Navon, I. Steinfeld, D. Lipson, Z. Yakhini, GOrilla: a tool for discovery and visualization of enriched GO terms in ranked gene lists. BMC Bioinformatics 10, 48 (2009).

