## Supplemental Note for "Detecting selection with a genetic cross"

**Trait-based tests of selection**

A comprehensive review of trait-based tests of selection can be found in Chapter 12 of Walsh and Lynch (1) (pp 438-478). Here I summarize the aspects most relevant to this work.

Trait-based selection tests (also referred to as tests of neutrality, since they are all based on null models of neutral evolution) can be broadly grouped into four major classes. These are: rate tests, time series tests, Q_ST_/driftsel, and QTL/GWAS-based tests.

As described in the main text, rate tests (such as Lande’s F_CV_ and F_MDE_) are considered to be qualitative guides rather than statistical tests (1, 2). Some of the assumptions made by these tests include that N_e_ (effective population size) and V_A_ (additive phenotypic variance) are both constant since the divergence of all lineages being tested, and that the mutational variance $\sigma_{m}^{2}$ measured in the lab (typically under very small N_e_) is representative of mutational variance in the wild, where the effectively neutral fraction of mutations is likely to be much smaller and could also be environmentally dependent. Considering these assumptions together with the high degree of uncertainty in estimating these parameters and their sampling variances, the results from these rate tests should be interpreted with caution.

In Q_ST_ methods, Q_ST_ > F_ST_ (where F_ST_ is genetic differentiation between populations) suggests directional selection, though there are several caveats associated with this. First, the absence of epistasis and dominance is assumed (though this may not greatly affect the results for most traits, particularly since most genetic variation in quantitative traits is additive (3)). Second, F_ST_ estimates are assumed to be based on neutral loci with the same mutational structure (e.g. mutation rate and propensity for back-mutations) as the loci affecting the trait of interest (4); it is generally impossible to know if this assumption is satisfied. Third, the use of full sibs to estimate Q_ST_ conflates additive genetic variation with dominance and maternal effects. Several reviews discuss these issues further (1, 5, 6), while others have taken a taken a more positive view (7–9).

Another class is QTL and GWAS-based tests in which a trait is first mapped to polymorphic loci and then phenotypic effects of these loci are compared to a neutral null model. The most widely used of these is the QTL sign test discussed in the main text, whose robustness is due to the minimal assumptions of its null model. The only assumption whose violation would lead to false positives in the sign test is that of QTL independence, where two or more QTL were in fact caused by the same QTN. Other factors can affect the sign test’s power, but will not lead to false positives. For example, the sign test has maximal power when QTL mapping is done optimally, but this is not an assumption of the method since sub-optimal QTL designs (e.g. small sample size or not raising all individuals in the same environment) will only reduce the signal. Likewise, although the sign test ignores epistasis between QTL, the presence of epistasis will not bias the test towards false positives. This dearth of assumptions makes the sign test’s null model the most robust of any test discussed here, though it pays for this robustness with power. As stated in the main text, at least 8 QTL must be mapped to reach a nominal p < 0.01 (observing all 8 in the same direction is only expected in 1 out of 2^7^ neutral traits; it is not 2^8^ because there are two possible directions in which the QTL could all act, i.e. the test is two-sided). If one QTL goes in the minority direction, then 12 QTL are required to reach p < 0.01. Few QTL studies reach even 8 QTL per trait (e.g. see Rieseberg (10) Supp Tables 4-5; note that most of the traits in these tables with >8 QTL are actually from combining QTL measured in different environments. For example, the 23 QTL listed for fruit fly lifespan in that paper’s Supp Table 5 were measured in five environments (11). Some of these QTL overlap and may be caused by the same QTN, so for the purpose of the sign test the relevant number of QTL is per environment tested, which is rarely >8).

Several methods that have been claimed to measure selection on quantitative traits such as gene expression were not included in this discussion because they cannot actually detect selection. For example, the ratio of a gene’s expression level variance between species to its variance within species (superficially analogous to Equation 1) has been used as a metric of selection, but since it is not known how neutral traits would be distributed for this metric, it cannot distinguish neutrality from selection (12). In addition, the correlation between a gene’s expression level and fecundity (while controlling for other gene expression levels) has been used to investigate selection acting on gene expression (13). This assumes that causality only goes from expression to fecundity; but we know that other factors (such as overall health) affect both gene expression and fecundity, so any observed correlations do not imply causality. As an analogy, this method is equivalent to claiming that any gene differentially expressed in individuals with a disease (after controlling for other gene expression levels) must be causing that disease.

As described in the main text, a caveat shared by all of these tests is trait ascertainment bias. In some cases it is possible to correct for this, either by modifying the test itself or by using a more conservative p-value threshold. For example, if an investigator chooses to test only the most divergent trait out of 100 possible candidates, then a Bonferroni correction for 100 tests would be a conservative correction (since this assumes that all traits are independent, which may not be the case). In many cases the number of possible candidate traits may be difficult to estimate, in which case the investigator could state that a particular trait rejects the neutral null model as long as it was selected from fewer than 0.05/p candidate traits, where p = the p-value of the selection test for that trait. In the case of the QTL sign test, one can condition on parental trait divergence to control for ascertainment (14). In the case of the *v*-test, conditioning on parental divergence is not possible since this would be conditioning away a key input for the test.

One final note about all trait-based tests of selection is that they cannot reveal what traits were actually under selection; rather they test what traits have been *affected by* selection. This could act via selection on the trait itself, or via pleiotropic effects of another trait. This issue is largely semantic, depending on how one defines a trait.

**The *v*-test equation**

Equation 2 differs from Equation 1 in two respects: a correction for when H^2^ < 1, and a constant term. The goal of the H^2^ correction is to include only the genetic fraction of phenotypic variance. For $\sigma_{F2}^{2}$ this is achieved by multiplying by H^2^. For $\sigma_{par}^{2}$ it is different since this is a variance of parental strain means. In this case the variance of each parental strain’s sample mean is subtracted from $\sigma_{par}^{2}$ to remove any environmental variance. Alternatively, the among-parent phenotypic variance (the numerator of Equation 2) could be estimated using one-way ANOVA (Equation 12.8e of Walsh and Lynch (1)). The broad-sense heritability H^2^ is used in Equation 2 rather than narrow-sense h^2^ because H^2^ includes non-additive (dominant and epistatic) variance, which if present will contribute to $\sigma_{par}^{2}$; to remove this only from the denominator by using h^2^ would lead to an unfair comparison (see “Possible extensions to the *v*-test” below for further discussion).

The constant c in Equation 2 reflects the need for the mean of *v* to equal one. Any factors that shift the distribution of cross progeny phenotypic variance relative to parental variance can be corrected with this constant (changes in the distribution of QTL effect sizes do not need to be corrected because they tend to affect both variance terms equally under neutrality). The value of the constant is equal to the ratio of expected phenotypic variances in the parents divided by that of the cross progeny. A convenient way to calculate this is based on a single locus that leads to a parental phenotypic difference of 2. The parental values will be [0 2] (the baseline value is set to zero without loss of generality) and thus $\sigma_{par}^{2}$ = 1 (which is the population variance rather than the sample variance). This will always be the numerator of c.

The denominator depends on the type of cross and dominance. For example, in an F_2_ cross between inbred parents with genotypes AA and BB there are four possible genotypes at each locus: AA, AB, BA, and BB. If each B allele adds 1 to the trait value (i.e. no dominance) then the four trait values are [0 1 1 2]. The variance is ½, so in this case c = 2. If instead all loci are dominant in the same direction (unidirectional) then the same genotypes yield a phenotype vector [0 2 2 2] and c = 4/3. If all loci are dominant but with 50% in each direction (bidirectional), the phenotypes are [0 0 2 2] and c = 1 (in this case the heterozygote 0 and 2 do not correspond to AB and BA genotypes, but rather reflect the fact that half the heterozygotes will have a phenotypic value 0 and the other half will have 2). Recombinant inbred lines have genotypes AA and BB, phenotypes [0 2], and c=1 (regardless of dominance). In the case of a backcross to the AA parent, the possible genotypes are AA and AB; under additivity the resulting phenotypes are [0 1] and c = 4, while if B is always dominant then the phenotypes are [0 2] and c = 1. For an F_3_ cross the genotype frequencies are 3/8 AA, 1/8 AB, 1/8 BA, and 3/8 BB, so the phenotype vector under additivity is [0 0 0 1 1 2 2 2] and c = 4/3. For haploids the possible genotypes are A and B, the phenotype vector is [0 2], and c = 1 (hence the lack of this constant in Equation 1).

This approach can be adapted for any scenario where the expected phenotype vector can be defined. This includes cases of partial dominance, overdominance, segregation distortion, etc. To account for a mixture of different dominance effects across loci, the phenotype vector should include the expected phenotype values in their expected proportions. For example, in an F_2_ cross with half additive and half bidirectional dominant loci the phenotypes would be [0 1 1 2 0 0 2 2] and c = 4/3.

In practice, dominance can be estimated by partitioning the phenotypic variance among cross progeny (15). In some cases it is possible to infer dominance in the F_1_ (when the F_1_ phenotype matches one of the parents, suggesting unidirectional dominance), but subsequent generations provide more information. Since additive effects always call for a larger value of c than dominance (for a given cross design), which will decrease *v*, if a particular trait’s dominance is not known then one should conservatively (from the perspective of detecting directional selection) assume additivity.

Note that in cases when zero phenotypic variation is expected in the cross progeny (e.g. a backcross to a fully dominant parent), c is infinite, so the test cannot be performed.

**The *v*-test in practice**

In general, any factors that inflate *v* above its true value will lead to underestimation of p_nut_, whereas factors that decrease *v* below its true value will lead to overestimation. For example:

- Environment: all individuals should be raised in the same environment. Random environmental variance will decrease H^2^, the genetic component of phenotypic variance. If H^2^ can be estimated accurately, this will have little impact on test performance; if environmental variance is not accounted for, *v* will be more constrained around its mean of one (as in Fig 2d), decreasing the power of the test to reject the null in either direction.
- Inheritance: all loci should be inherited equally from each parent. For autosomal loci this is generally true except in cases of segregation distortion. It may be possible to correct for such distortion, either through the constant in Equation 2 (see below) or by genotyping individuals and excluding some to ensure equal contribution of each parent. For sex chromosomes, mitochondrial genomes, and maternal effects, equal representation can be ensured with reciprocal crosses. Alternatively, if these only make a minor contribution to the trait of interest then their effects can be ignored.
- Homozygosity: if parental lines are heterozygous at QTN affecting the trait of interest, then the F_1_ generation will have a mix of homozygous and heterozygous individuals at these loci. This is not an issue if the F_2_ generation is derived from an equal number of both types of F_1_, but if not, parental heterozygosity could either increase *v* (if the F_1_ is mostly homozygous) or decrease *v* (if the F_1_ is mostly heterozygous). For most crosses this is not relevant since the parents are inbred and/or highly divergent (thus likely to be homozygous for most QTN), and the F_1_ individuals used are unlikely to be biased in their heterozygosity at any loci.
- Polygenicity: the *v*-test is designed for polygenic traits with many QTN segregating independently in the cross progeny. Reducing the number of independently segregating QTN (or equivalently, the recombination index; Fig 2b) below ~20 tends to constrain the distribution of F_2_ trait values (resulting in a tighter distribution around its expectation of half the parental variance in diploids, which in turn leads to a tighter distribution of *v* around 1) and thus decrease the power of the test to reject the null in either direction.
- Heritability: increasing the H^2^ value in Equation 2 leads to lower values of *v*. Therefore if H^2^ cannot be accurately estimated, a conservative approach (from the standpoint of detecting directional selection) is to use a value larger than what is likely to be true. Alternatively, one can conclude that the null hypothesis can be rejected as long as H^2^ is less than some value. The *v*-test is two-sided, with low values of *v* indicating stabilizing selection (Fig 1c), but the power to detect directional selection is more robust to low H^2^ than is the power to detect stabilizing selection. Because $\sigma_{par}^{2}$ is bounded at zero, and its null distribution has its highest density near zero, there is only a narrow range of $\sigma_{par}^{2}$ values that are significantly less than expected; thus even a small amount of noise (e.g. environmental or technical) in parental phenotype data could push the variance outside this range. In contrast, the same level of noise in phenotypically divergent parents would have a smaller proportional effect on $\sigma_{par}^{2}$, and thus less impact on the power to detect directional selection.
- Additivity: similar to many quantitative genetic methods, the *v*-test assumes that QTN are additive. If instead they are multiplicative (e.g. a QTN increases size by 10% rather than 10 cm), then a log transformation is warranted. Additional transformations to achieve additivity are also possible (16).
- Normality: the *v*-test assumes the trait follows an approximate normal distribution in the F_2_ (or other cross) populations. For a trait that is not normally distributed, a transformation may render it additive (16). However if it is multimodal, this is more likely due to a very small number of loci explaining the trait (e.g. a monogenic trait will be trimodal in the F_2_), in which case the test has little power.
- Biased cross progeny: If some cross progeny are not observed (e.g. due to embryonic lethality), and this affects the phenotypic variance among the observed progeny, this will bias the test. However, this will only be an issue in crosses with strong genetic incompatibilities linked to QTN that affect the trait of interest. If lethality is not complete among the progeny with less fit genotypes, then its effects could be mitigated by genotyping and then selecting progeny with no genotypic bias.
- Quantitative traits: The test is designed for quantitative traits that can have many possible values. It has very little power for binary traits, since as long as the F_2_ mean trait value is equal to the midparental value, $\sigma_{par}^{2}$ will equal $\sigma_{F2}^{2}$ (even with hundreds of reinforcing QTN).

These points can be summarized into a set of “best practices” for *v*-test experimental design. The power of the *v*-test is greatest for RILs (due to their higher effective RI and also the lack of any need to account for dominance effects). Polygenic traits are ideal, though traits of unknown genetic architecture are acceptable since monogenic traits will result in conservative p-values rather than false positives. Species with high RI will also yield greater power, though the effect is small at RI > ~20. Crosses with segregation distortion or lethality due to incompatible genetic interactions should be avoided. Reciprocal crosses are ideal if there is a significant contribution from sex chromosomes, the mitochondrial genome, or maternal effects. Parents that are homozygous for all major QTN are preferred. Traits should be tested for additivity and normality, and transformed if necessary. To maximize power when only a fixed number of individuals can be phenotyped, in general the number of progeny should be roughly equal to the number of parents phenotyped. However, to detect stabilizing selection a modest excess of parental individuals is ideal (due to the sensitivity to noise when the parental phenotypes have little divergence; see “Heritability” point above), while to detect directional selection a modest excess of progeny is recommended. If there is an *a priori* hypothesis of either directional or stabilizing selection then a one-sided test is appropriate; otherwise a two-sided test should be used. $\sigma_{par}^{2}$ should always be calculated as a population variance (since the two parents are the complete set of parents), whereas $\sigma_{F2}^{2}$ should be a sample variance (since all possible F_2_ progeny have not been sampled).

**The Castle-Wright estimator**

The CW estimator estimates the number of loci underlying a trait’s divergence between two parental lines, assuming that loci act additively, are unlinked, have equal effect sizes, and have fully reinforcing directionalities in the parents (as in Fig 1a right). Since at least some of these assumptions are always violated in real data, the result is referred to as the “effective” number of loci (*n*), which is always an underestimate of the true number (15). As expressed by Wright (Equation 15.8) (16),

$$n=\frac{{(\bar{P}_{2}-\bar{P}_{1})}^{2}}{8(\sigma_{F2}^{2}-\sigma_{E}^{2})}$$

where $\bar{P}_{2}$and $\bar{P}_{1}$ are the parental strain means, $\sigma_{E}^{2}$ is the phenotypic variance due to environment, and $\sigma_{F2}^{2}$ is the phenotypic variance in the F_2_. The $\sigma_{F2}^{2}-\sigma_{E}^{2}$ term is also known as the segregation variance $\sigma_{S}^{2}$, which is equal to $\sigma_{F2}^{2}H^{2}$. Cockerham introduced a correction for the effect of environmental variation on the parental means (17):

$$n=\frac{{(\bar{P}_{2}-\bar{P}_{1})}^{2}-\sigma_{\mu1}^{2}-\sigma_{\mu2}^{2}}{8\sigma_{S}^{2}}$$

where the sampling variance of the parental means $\sigma_{\mu1}^{2}=\sigma_{p1}^{2}/n_{p1}$ and $\sigma_{\mu2}^{2}=\sigma_{p2}^{2}/n_{p2}$.

Based on the fact that $\sigma_{par}^{2}={(\bar{P}_{2}-\bar{P}_{1})}^{2}/4$, it can be seen that Cockerham’s formulation is identical to Equation 2 when no dominance is present (i.e. c = 2).

Cockerham also pointed out that $\sigma_{S}^{2}$ is ideally estimated by least squares when data are available for both parents, the F_1_, F_2_, and both backcrosses (17). This reasoning applies to estimating *v* as well, though this approach was not taken here since F_2_ data are far more widely available than matched backcross+F_2_ data.

Several improved estimators of *n* have also been proposed (18, 19). However these do not perform well in neutral simulations (data not shown), since the actual number of loci underlying a trait (the quantity that *n* aims to estimate) is not equivalent to, or even correlated with, *v* (as discussed in the main text).

As mentioned in the main text, the equivalence of the CW estimator with Equation 2 means that published CW estimator values can be used directly as tests of neutrality. However, care must be taken to know which version of the CW estimator was used and what assumptions it makes. For instance, most published applications of the CW estimator do not use Cockerham’s modification, so if one wishes to reinterpret these CW estimates as estimates of *v*, the variance of the parental means should be much greater than the sum of their sampling variances.

As an example I will use six traits measured in F_2_ crosses and backcrosses for all 12 possible reciprocal pairings among four species of *Mimulus* (monkeyflower) (20). Two of these species are selfers and two are outcrossers, and the six traits are all related to reproductive timing or morphology. CW estimators were calculated with Cockerham’s modification, and the authors additionally estimated dominance for each trait and accounted for this in their CW estimates. Examining the CW estimator values in their Table 3, several trends are apparent. Mean CW values are all in the range of 5.3-12.8 for the five morphological traits (corolla width, corolla length, stamen level, pistil length, and stigma-anther separation; corresponding to 5x10^-4^ < *v*-test p_nut_ < 0.024) in the four crosses between selfers and outcrossers (see the bottom row in their Table 3 for the means across these four crosses); in contrast, the CW values for these traits between the two selfers are much lower, between 0.2-1.4 (corresponding to 0.24 < *v*-test p_nut_ < 0.66). This difference suggests the strength of selection on these morphological traits differs between selfers and outcrossers. In contrast, the only non-morphological trait (flowering time) showed the opposite trend, with greater CW estimates in the cross between selfers (5.1, *v*-test p_nut_ = 0.026) than between selfers x outcrossers (mean = 0.8, *v*-test p_nut_ = 0.37). Therefore flowering time shows no difference in selection strength between selfers and outcrossers, and seems to be subject to very different patterns of natural selection as compared to the five morphological traits. In sum, the reinterpretation of CW estimates as tests of neutrality allow inferences about selection to be made even when the phenotype data are not available.

**Epistasis and selection**

Epistasis (when the phenotypic effects of a variant depend on genomic background) can profoundly affect phenotypic variation. In the main text, simulations are focused on broad-scale patterns like diminishing returns epistasis. Exploration of other transformations of trait values to mimic other types of epistasis showed little impact, as in Fig 2e. For example, an additive model with a strict maximum trait value (set at the median F_2_ value, so half of all F_2_ individuals and one parent was assigned the median F_2_ value) still adhered to the expected distribution (not shown). Epistasis affecting the population mean (e.g. heterosis) is not relevant to the *v*-test, since *v* depends only on trait variances, which should be independent of the mean (if they are not, then a transformation is warranted (16)).

Extra scrutiny is needed for the case where epistatic interactions in the parents have been shaped by selection. To understand this at an intuitive level, it helps to distinguish between epistasis that is random with respect to parental divergence vs. non-random (i.e. systematically increasing or decreasing parental divergence more than expected by chance). Both types will result in similar effects on F_2_ individuals (and thus $\sigma_{F2}^{2}$), since the parental directionalities are randomized in the F_2_. Note that this classification is independent of concepts such as positive/negative epistasis and increasing/decreasing returns epistasis.

An analogy can be drawn with Fig 1a, where instead of each row being an individual QTL, it is a combination of two or more loci that interact epistatically. Random epistasis is the situation in Fig 1a left, while non-random (due to directional selection) is on the right. Since random epistasis by definition has no directionality bias in either parents or F_2_, it tends to increase both $\sigma_{F2}^{2}$ and $\sigma_{par}^{2}$ but not their ratio. Therefore the effects of epistasis on *v* are modest as long as most heritable phenotypic variation is additive, as is usually the case for complex traits (3). However, if random epistasis is strong enough that it dominates the trait variation, then both $\sigma_{F2}^{2}$ and $\sigma_{par}^{2}$ will be dominated by randomness and the *v*-test will tend to be conservative, having less power to reject the null in either direction. Mathematically this is similar to Fig 2d left panels, where random noise was added to trait values to simulate environmental variation; sufficiently complex epistasis would also resemble random noise, even if it is not truly random.

If instead epistasis is non-random, this means that its parental directionality is systematically biased in one direction. Using the same logic as the sign test in Fig 1a, it is clear that this non-random epistasis must be due to selection rather than neutral evolution; it reflects co-adapted alleles (also known as co-adapted gene complexes) in each parent. These co-adapted alleles could either increase parental divergence under directional selection, or decrease it under stabilizing selection. In either case, if selection is involved then ideally a test of selection would include its effects in its assessment of the neutral null hypothesis. The parental variance (Equation 2 numerator) includes any effects of epistasis on parental traits, so it is naturally incorporated into estimates of *v*.

Put another way, the logic of using F_2_ progeny as the neutral null model for parental traits applies equally to additive or epistatic variation, so no modification to Equation 2 is necessary to account for epistasis. In fact, one could imagine a version of the *v*-test focused exclusively on epistasis, comparing the trait variance due to epistasis in parents vs. F_2_ progeny. Deviation of parental epistatic variance from that in the F_2_ would reject the null hypothesis of neutrality.

**Notes on specific traits under selection**

*Head shape in Hawaiian* Drosophila

The trait with strongest evidence of directional selection—even stronger than any artificially selected trait in Fig 4a—was head shape in a cross between two fruit fly species, *D. silvestris* and *D. heteroneura.* The males of *D. heteroneura* (and to a lesser extent the females) have extremely wide heads, which females of this species prefer when selecting mates (21). For a summary of the debate surrounding the role of sexual selection in this trait, see Boake (21).

The results in Supp Table 1 were calculated by combining F_2_ individuals from all crosses reported by Templeton (22), separately for males and females. However, head shape is quite different in each direction of reciprocal F_1_ crosses, suggesting potential importance of separating these crosses for analysis. Performing the *v*-test on each F_2_ cross separately revealed significant effects for each (p_nut_ = 4.2x10^-9^ and 2.8x10^-4^). An F_6_ cross (performed in only one direction) further validated these results (p_nut_ = 1.6x10^-4^).

Templeton noted that head size and shape both vary in these flies, so to focus on shape only, he transformed all measurements to achieve the same head radius but preserve shape (22). After this transformation, head length had a smaller $\sigma_{F2}^{2}$, which is why it is the more significant trait in Supp Table 1. However, head width is more divergent between the species. Therefore it may be best to consider this trait as head shape, rather than specifically as width or length.

*Human skin color*

Harrison and Owen (23) identified people—mostly residents of Liverpool, England—whose ancestry was either West African, European, or a combination of both. Those individuals for whom admixture occurred within the last two generations could be classified as “F_1_”, “F_2_”, or “backcross”. Skin color was then measured with a reflectance spectrophotometer. The authors assessed the fit to an additive model at each wavelength, and arrived at three acceptably additive wavelengths, each requiring a different transformation (log, antilog, and no transformation). These three transformed wavelengths are what was used in this study.

*Burrowing behavior in* Peromyscus

Weber et al. (24) performed a backcross between two species of *Peromyscus* mice, *P. maniculatis* and *P. polionotus*. In addition to the 33 parental mice phenotyped in the original study, data for 21 more were kindly provided by Dr. Caroline Hu. Several different subsets of burrow length were measured (e.g. entrance tunnel length, escape tunnel length), both as a mean value for each mouse across multiple trials and as a maximum value per mouse. Burrowing behavior showed nearly complete dominance in the F_1_ and the backcross (which was close to the midparent value as expected for a dominant trait), so c was set to 1 (see above). The log transformation was used to achieve additivity. The most significant trait was mean total burrow length, and other correlated traits were also significant (Supp Table 1). The H^2^ estimate (see Methods) for this trait was 0.13 (resulting in p_nut_ = 1.1x10^-7^), but since the parental mice were not inbred, within-strain variance may include a genetic component and thus lead to underestimation of H^2^. Therefore in Supp Table 1 I conservatively set H^2^ = 0.4 (resulting in p_nut_ = 2.1x10^-3^), which is higher than the heritability of most behavioral traits (25). The trait remains significant at higher heritabilities as well (e.g. p_nut_ = 0.012 at H^2^ = 0.6).

**Notes on Fig 4d**

Comparing p_nut_ values across species requires attention to potential artifacts. For example, sample size and RI can affect *v* (Fig 2b-c), raising the question of whether they could affect the results in Fig 4d. However there was no association between the stabilizing selection metric and these parameters in Fig 4d; for instance, out of the six species yeast has both the largest sample size (n_F2_ = 1012) and highest RI (~73), but had the most neutral-like p_nut_ distribution, which is the opposite of the expectation if these parameters influenced the stabilizing selection metric.

A second question is whether the π of each species could affect the *v*-test’s power to detect stabilizing selection, rather than the strength of selection itself. Species with higher π may have more segregating QTN due to their higher overall genetic variation, and thus potentially more power to reject neutrality. However, as discussed in the main text, a more important determinant of power is RI; even a species with many QTN will have little power in the *v*-test if these QTN remain linked due to a low RI (Fig 2b). Since RI is not associated with stabilizing selection in these species (as mentioned above), π is unlikely to be a major factor in determining power. Higher π values also suggest a longer coalescent time (time to the most recent common ancestor); however this should not affect the test aside from the effect on QTN mentioned above, since the only input for the test is trait values determined by the QTN.

It is possible that each species’ heterozygosity could affect the measurement of gene expression levels through a technical artifact: either the mappability of RNA-seq reads (for the yeast data) or the suboptimal hybridization of mismatched microarray probes (for the other five species). This would add to the variance of each affected mRNA level in the same way as a cis-eQTL, since it would depend on the local genotype. Like for a QTL, the effect on $\sigma_{F2}^{2}$ depends only on the effect size. However, the effect on $\sigma_{par}^{2}$ (and *v*) also depends on directionality: for a given gene it increases (decreases) $\sigma_{par}^{2}$ if the parent with higher mRNA level has less (greater) divergence from the reference sequence for that gene. To test for this potential artifact, I asked whether cis-eQTL in *B. rapa* (the species with the highest π and strongest skew towards stabilizing selection; Fig 4d right panel) show any bias in their directionality, as would be expected if local genotype were affecting probe hybridization and decreasing apparent expression preferentially for alleles from the strain with more mismatches. There was an almost perfect balance of directionality (50.2% of the 3303 cis-eQTL (26) increased expression of the IMB211 allele in the normal phosphorus condition, and 49.6% in low phosphorus). Therefore no bias is apparent even in the species with the greatest potential for such a bias.

Furthermore, even if this bias were present it would be unlikely to lead to the relationship observed in Fig 4d. To cause this relationship, the bias would have to preferentially reduce $\sigma_{par}^{2}$ for thousands of genes in species with higher π. Note that since overall expression levels are always normalized to be similar or identical between strains, one parent rarely has higher expression for much more than half the genes; therefore this pattern could not be caused by a strain with higher overall mRNA levels also being more divergent from the reference sequence genome-wide, since this would be canceled out by a nearly equal number of genes acting in the opposite direction. The relationship in Fig 4d could only be caused at an individual gene level, with parent A more divergent from the reference sequence specifically in those genes where it also has higher expression than parent B. This scenario seems unlikely to occur at a high enough frequency to cause the shift in p_nut_ values seen from yeast to *B. rapa* (Fig 4d, compare left panel to right panel), since 1) there no obvious mechanism to link up-regulation of individual genes with increased divergence from the reference genome, and 2) there is no directionality bias for cis-eQTL even in *B. rapa*, the species with highest π.

**Possible extensions to the *v*-test**

- Correcting for RI: As discussed in the main text and above, RI < ~20 constrains the values of *v* to be more tightly distributed around 1, and thus reduces power to reject the null in either direction. Since RI is known for almost every species that is used for genetic crosses, a modification to Equation 2 can be envisioned to correct for this conservativeness.
- Confidence intervals: When *v* is based on a very small sample size (e.g. n_F2_ = 3; Fig 2c) it has a large variance that can lead to either over- or under-estimation. Confidence intervals could be estimated for *v*, which could aid in its interpretation. This has been derived analytically for the equivalent CW estimator (Equation 9.25 of Lynch and Walsh (15)) or could be based on bootstrapping, though larger sample sizes would be advisable for the latter.
- Multiple traits: When multiple traits are measured for the same cross, it may be possible to increase the power of the *v*-test by estimating their genetic covariance and considering them jointly (27, 28).
- Additive effects: In some cases it may be desirable to test only the additive genetic variance for selection, since selection is most effective on additive effects (due to being passed on without being shuffled in each generation like epistatic effects). This could be achieved with a modification to Equation 2, replacing H^2^ in the denominator with h^2^, and replacing $\sigma_{par}^{2}$ with $\sigma_{EBV}^{2}$, which is the variance of the estimated breeding values for the two parents; $\sigma_{p1}^{2}$ and $\sigma_{p2}^{2}$ would likewise be replaced by the within-strain variance in EBVs for each parental strain.
- Admixed populations: Hybridization between populations and even species is widespread in nature. The *v*-test could be adapted to work in any admixed population if the trait is measured in both parental populations and admixed individuals, and the admixture proportions in every phenotyped individual are known. Then the constant c in Equation 2 (discussed above) could be estimated using the same general approach: expected parental variance divided by expected admixed progeny variance. However, three assumptions are inherent in this. First, population-level admixture proportions are assumed to be equal throughout the genome (or more pertinently, at all loci affecting the trait). Second, any correlation between environment and admixture proportion could bias the test. Third, if selection among admixed individuals affects phenotypic variance, the hybrid phenotypes will not represent what would have been generated by chance (see “Biased cross progeny” above). However this does not violate the neutral null model, since this selection would not happen under neutral evolution.

**Supplemental References**

1.	B. Walsh, M. Lynch, *Evolution and Selection of Quantitative Traits* (2018).

2.	M. Turelli, J. H. Gillespie, R. Lande, RATE TESTS FOR SELECTION ON QUANTITATIVE CHARACTERS DURING MACROEVOLUTION AND MICROEVOLUTION. *Evolution (N. Y).* **42**, 1085–1089 (1988).

3.	W. G. Hill, M. E. Goddard, P. M. Visscher, Data and Theory Point to Mainly Additive Genetic Variance for Complex Traits. *PLoS Genet.* **4**, e1000008 (2008).

4.	Z. Li, A. Löytynoja, A. Fraimout, J. Merilä, Effects of marker type and filtering criteria on QST-FST comparisons. *R. Soc. Open Sci.* **6** (2019).

5.	B. PUJOL, A. J. WILSON, R. I. C. ROSS, J. R. PANNELL, Are Q ST - F ST comparisons for natural populations meaningful? *Mol. Ecol.* **17**, 4782–4785 (2008).

6.	M. C. WHITLOCK, Evolutionary inference from Q ST. *Mol. Ecol.* **17**, 1885–1896 (2008).

7.	J. Merilä, P. Crnokrak, J. Merilä, Comparison of genetic differentiation at marker loci and quantitative traits. *J. Evol. Biol.* **14**, 501 (2001).

8.	T. Leinonen, R. B. O’Hara, J. M. Cano, J. Merilä, Comparative studies of quantitative trait and neutral marker divergence: A meta-analysis. *J. Evol. Biol.* **21**, 1–17 (2008).

9.	J. K. McKay, R. G. Latta, Adaptive population divergence: markers, QTL and traits. *Trends Ecol. Evol.* **17**, 285–291 (2002).

10.	L. H. Rieseberg, A. Widmer, A. M. Arntz, J. M. Burke, Directional selection is the primary cause of phenotypic diversification. *Proc. Natl. Acad. Sci.* **99**, 12242–12245 (2002).

11.	C. Vieira, *et al.*, Genotype-environment interaction for quantitative trait loci affecting life span in Drosophila melanogaster. *Genetics* **154**, 213–227 (2000).

12.	H. B. Fraser, Genome-wide approaches to the study of adaptive gene expression evolution. *BioEssays* **33**, 469–477 (2011).

13.	S. C. Groen, *et al.*, The strength and pattern of natural selection on gene expression in rice. *Nature* **578**, 572–576 (2020).

14.	H. A. Orr, Testing natural selection vs. genetic drift in phenotypic evolution using quantitative trait locus data. *Genetics* **149**, 2099–104 (1998).

15.	M. Lynch, B. Walsh, *Genetics and analysis of quantitative traits* (1998).

16.	S. Wright, *Evolution and the Genetics of Populations, Volume 1* (1968).

17.	C. C. Cockerham, Modifications in estimating the number of genes for a quantitative character. *Genetics* **114**, 659–664 (1986).

18.	Z. B. Zeng, Correcting the bias of Wright’s estimates of the number of genes affecting a quantitative character: a further improved method. *Genetics* **131**, 987–1001 (1992).

19.	S. P. Otto, C. D. Jones, Detecting the undetected: estimating the total number of loci underlying a quantitative trait. *Genetics* **156**, 2093–107 (2000).

20.	C. B. Fenster, K. Ritland, Quantitative genetics of mating system divergence in the yellow monkeyflower species complex. *Heredity (Edinb).* **73**, 422–435 (1994).

21.	C. R. B. Boake, Sexual selection and speciation in Hawaiian Drosophila. *Behav. Genet.* **35**, 297–303 (2005).

22.	A. R. Templeton, Analysis of Head Shape Differences Between Two Interfertile Species of Hawaiian Drosophila. *Evolution (N. Y).* **31**, 630 (1977).

23.	G. A. HARRISON, J. J. OWEN, STUDIES ON THE INHERITANCE OF HUMAN SKIN COLOUR. *Ann. Hum. Genet.* **28**, 27–37 (1964).

24.	J. N. Weber, B. K. Peterson, H. E. Hoekstra, Discrete genetic modules are responsible for complex burrow evolution in Peromyscus mice. *Nature* **493**, 402–405 (2013).

25.	D. G. Stirling, D. Réale, D. A. Roff, Selection, structure and the heritability of behaviour. *J. Evol. Biol.* **15**, 277–289 (2002).

26.	J. P. Hammond, *et al.*, Regulatory Hotspots Are Associated with Plant Gene Expression under Varying Soil Phosphorus Supply in Brassica rapa. *Plant Physiol.* **156**, 1230–1241 (2011).

27.	A. Kremer, A. Zanetto, A. Ducousso, Multilocus and multitrait measures of differentiation for gene markers and phenotypic traits. *Genetics* **145**, 1229–1241 (1997).

28.	O. Ovaskainen, M. Karhunen, C. Zheng, J. M. C. Arias, J. Merilä, A new method to uncover signatures of divergent and stabilizing selection in quantitative traits. *Genetics* **189**, 621–632 (2011).
